# DeepHINT: Understanding HIV-1 integration via deep learning with attention

**DOI:** 10.1101/258152

**Authors:** Hailin Hu, An Xiao, Sai Zhang, Yangyang Li, Xuanling Shi, Tao Jiang, Linqi Zhang, Lei Zhang, Jianyang Zeng

## Abstract

**Motivation:** Human immunodeficiency virus type 1 (HIV-1) genome integration is closely related to clinical latency and viral rebound. In addition to human DNA sequences that directly interact with the integration machinery, the selection of HIV integration sites has also been shown to depend on the heterogeneous genomic context around a large region, which greatly hinders the prediction and mechanistic studies of HIV integration.

**Results:** We have developed an attention-based deep learning framework, named DeepHINT, to simultaneously provide accurate prediction of HIV integration sites and mechanistic explanations of the detected sites. Extensive tests on a high-density HIV integration site dataset showed that DeepHINT can outperform conventional modeling strategies by automatically learning the genomic context of HIV integration solely from primary DNA sequence information. Systematic analyses on diverse known factors of HIV integration further validated the biological relevance of the prediction result. More importantly, in-depth analyses of the attention values output by DeepHINT revealed intriguing mechanistic implications in the selection of HIV integration sites, including potential roles of several basic helix-loop-helix (bHLH) transcription factors and zinc-finger proteins. These results established DeepHINT as an effective and explainable deep learning framework for the prediction and mechanistic study of HIV integration.

**Availability:** DeepHINT is available as an open-source software and can be downloaded from https://github.com/nonnerdling/DeepHINT

## 1 Introduction

Integration of the HIV-1 genome to human genome is a crucial step in viral infection and replication cycle. Clinically, the integration of HIV is closely related to the formation of latent viral reservoir and the rebound of viral load when antiretroviral therapy (ART) is interrupted [1–3]. Furthermore, recent studies have also revealed that the integration of HIV provirus within specific genes can affect the persistence of infected cells [4, 5], indicating a more significant role of the selection of HIV integration site in disease progression.

Despite the long-lasting research efforts, the detailed mechanisms and functional implications of the selection of HIV integration sites still remains largely unclear [6]. In addition to the local sequence motifs of the human genome that directly interact with the DNA integrase [7], previous researches have also associated the preference of HIV integration events with various genomic landmarks, e.g., the binding of integrase cofactor LEDGF/p75 [8], actively transcribed genes [9], intron regions [10], chromatin accessibility [11], and nucleus-pore proximal regions [12]. To integrate diverse genomic features for predicting HIV integration sites, several computational methods have been proposed [13, 14]. However, these methods strongly rely on explicit feature engineering and input from various other experimental data, e.g., RNA-seq, ChIP-seq and DNase-seq data, which may not be universally available for all the integration prediction tasks. In addition, the resolution and the scope of feature engineering also limit the interpretation of mechanistic insight from these methods, leading to the insufficient usage of currently available large-scale HIV integration data [15].

The recent decade has witnessed the boom of deep learning. In computational biology, deep learning has become the state-of-the-art prediction methods in many applications, e.g., identification of nucleotide-protein binding and nucleotide modification sites [16–19], prediction of the functional effects of noncoding sequence variants [20, 21], cancer genomics [22], translation initiation and elongation modeling [23, 24] and drug discovery [25, 26]. On the other hand, despite the superior prediction performance, the explainability and the understanding of feature organizations of deep learning models often lag behind, which not only limits the applicability of deep learning techniques in exploring unknown cellular mechanisms and gaining insights, but also raises potential concerns of using a black box. One possible strategy to increase the explainability of deep learning models is the introduction of attention mechanisms, which are particularly designed to extract important regions of the input data by training an additional neural network that learns the relative importance of each input position from local features [27]. Thus, when applied to genomic sequence data, the introduction of attention mechanisms is expected to reveal important sequence positions that shape the prediction results from the deep learning framework and thus provide potentially important mechanistic insights about the observed genomic phenomenon [28–30].

Here, to address the HIV integration site prediction problem, we have developed an attention-based deep learning framework, named DeepHINT (Deep learning for HIV INtegraTion) (Fig. 1), for accurately predicting HIV integration sites by automatically extracting important sequence features and genomic positions solely from primary DNA sequences. Our work represents the *first* attempt to model the selection of HIV integration site by deep learning approaches. Extensive tests have shown DeepHINT can achieve an accurate prediction performance and outperforms the current state-of-the-art prediction methods that leverages way more experimental data. The biological relevance of the DeepHINT prediction results has also been validated by the associations with known genomic markers of HIV integration. More importantly, the information derived from the incorporated attention mechanism clearly indicates the relative importance of each position in the input genomic context of the predicted HIV integration sites, which thus helps us to explain both local and distal genetic features captured by the deep learning model. In particular, our prediction results highlight the potential roles of several basic helix-loop-helix (bHLH) transcription factors and zinc-finger proteins, which may expand our current understanding of HIV integration site selection. All these results have demonstrated the effectiveness of our deep learning based prediction approach and also provided useful insights to facilitate the mechanistic studies of the HIV integration process.

**Figure 1:**
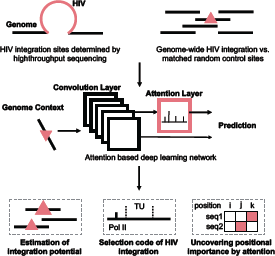
Schematic overview of the DeepHINT pipeline. In our prediction task, we used experimentally derived HIV integration sites as positive samples. To account for bias caused by enzyme digestion in the sequencing process, matched random control sites possessing the same distance distribution to the nearest enzyme digestion sites were generated as negative samples. See the main text for more details.

## 2 Methods

### 2.1 Designing an attention based deep learning architecture for modeling sequence context of HIV integration sites

#### 2.1.1 Feature Extraction by a convolutional neural network

DeepHINT first employs multiple convolution-pooling modules to automatically learn informative sequence features in the surrounding sequences of HIV integration sites (Fig. 2(a)). In particular, we first extend each HIV integration site both upstream and downstream by 1,000 bps to obtain the sequence context, resulting in a sequence profile denoted by *s* = (*nt*_1_, …, *nt*_2000_), where *nt_i_* stands for the nucleotide at the *i*th position. Each nucleotide in the sequence profile is then converted to a binary vector of length 4 by one-hot encoding, with each dimension corresponding to a nucleotide type. In the convolutional layer, a series of one-dimensional convolution operations are performed over the 4-channel input data, in which each channel corresponds to one dimension of the binary vector. In particular, each convolution operation corresponds to a weight matrix (i.e., kernel) that can also be regarded as a position weight matrix (PWM).

**Figure 2:**
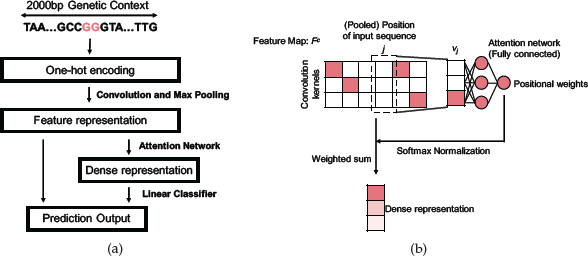
The deep learning framework implemented in DeepHINT. (a) The overall schematic view of the deep learning framework. (b) The illustration of attention mechanism. See the main text for more details.

More specifically, given a genomic sequence *s* = (*nt*_1_, …, *nt*_2000_) and the corresponding one-hot encoded representation *E*, the convolutional layer computes *X* = conv(*E*), i.e.,

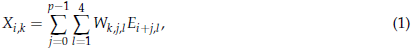

where 1 ≤ *i* ≤ 2000 *− p* + 1, 1 ≤ *k* ≤ *d*, *p* is the kernel size, *d* is the kernel number and *W* is the kernel weights. Next, we apply the rectified linear activation function (ReLU) on the convolution results, which mimics the biological neuron activation. After convolution and rectification, the max-pooling operators are used to perform dimension reduction. Therefore, through a series of convolution-pooling modules, the sequence profile can be compiled to a *d* × *q* feature map matrix (denoted by *F^c^*), where *q* represents the total (pooled) positions of the input sequence (Fig. 2(b)). As the kernel weights are trainable in the learning process, the convolution-pooling modules in our framework are expected to automatically extract important and diversified sequence features encoded in the genome context.

#### 2.1.2 Incorporation of the attention mechanism

To better capture and understand the positional importance of the sequence context, we further introduce an attention layer into our model (Fig. 2(a)). The attention layer takes the feature vector after convolution-pooling operations as input, and then computes a score indicating whether the neural network shall pay attention to the sequence features at that position. Basically, column *j* of the feature map matrix *F^c^* can be viewed as a feature vector (denoted by *v_j_*) that describes the feature of the *j*th position in the input sequence, with each dimension corresponding to a kernel in the convolution layer. The attention layer feeds each input feature to a shared feedforward neural network with a single hidden layer. The output of the attention layer is an importance score, denoted by *e_j_*, for which a larger value indicates that the corresponding position is more important for the contribution to final HIV integration site prediction. In particular, the columns of the feature map matrix *F^c^* are further averaged by taking the normalized importance scores *α_j_* as weights, resulting in a dense feature representation *F^a^*, i.e.,

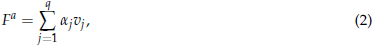

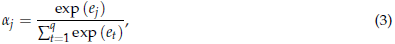

where *e_j_* is the importance score output by the shared neural network and *α_j_* is the corresponding normalized score.

To integrate the feature captured by the convolution-pooling modules (i.e., *F^c^*) and the attention mechanism (i.e, *F^a^*), we first concatenate all the values in *F^c^* matrix and linearly project them to one value (denoted by *S^c^*) that represents the contribution from a unified representation of the whole sequence. Finally, we concatenate *S^c^* with the dense representation *F^a^* and then feed them together to a logistic regression classifier to obtain a prediction score that indicates the probability of HIV integration. In summary, the full model can be expressed as

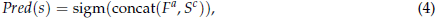

where *s* denotes the genomic context of a candidate integration site and

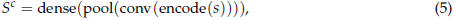

where encode(·), conv(·), pool(·), concat(·), dense(·) and sigm(·) represent the one-hot encoding, convolution, max pooling, concatenation, dense and sigmoid operations, respectively. Meanwhile, given a specific input sequence, we can also output a weight vector (denoted as *AttMap*)

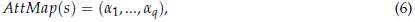

which expresses the model‘s attention on each position of the input sequence.

We note that in addition to mimicking the human intelligence in efficiently understanding images and texts with attention and deciphering the importance of individual features for prediction, the attention architecture also largely reduces the model complexity by globalizing the network structure and sharing weights in the neural network among the local feature vectors at different positions [27], which potentially overcomes the overfitting problem and thus leads to better model generalization.

### 2.2 Hyperparameter tuning and model training

Hyperparameter tuning in DeepHINT involves kernel size, the number of kernels and max-norm of weights of the convolution layer, the pooling length of the max-pooling layer, hidden layer size and activation function in the attention layer, dropout rate, and the optimizer algorithm. The calibration is performed with the tree-structured Parzen estimator (TPE) approach [31] using the Hyperas^1^ package. Specifically, 100 evaluations are carried out using separate training and validation sets, which includes the all positive samples and randomly selected negative samples with the same number. The best hyperparameter setting (Supplementary Table S1) is selected based on validation loss. The five-fold cross-validation test results for the calibrated hyperparameters are shown in Supplementary Figure S1.

After hyperparameter tuning, the deep neural network of DeepHINT is trained by minimizing the binary cross-entropy loss function, which is defined as the sum of negative log likelihood, i.e.,

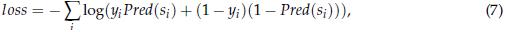

where *y_i_* stands for the true binary label of an input sequence *s_i_*. The standard error backpropagation algorithm [32] and the batch gradient descent method [33] are implemented for training. We also introduce several regularization techniques, including adding max-norm constraints on kernel weights [34], dropout [35] and early stopping [33] to alleviate the potential overfitting problem. In addition, to better address the data imbalance problem (negative samples are ten times as many as positive samples) and make full use of the excessive negative sample information, we also apply an bootstrapping-based training strategy [24, 36]. Specifically, in parallel, we train the deep neural network for 16 times with an equal number of randomly sampled (with replacement) positive and negative samples from the original training set. This strategy results in an ensemble of deep neural network classifiers, whose prediction scores and attention maps are averaged to give the final output, i.e.,

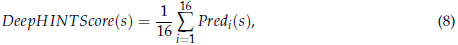

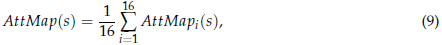

where *Pred_i_*(*s*) and *AttMap_i_*(*s*) represent the prediction score and the output attention map of a single deep neural network for a given input sequence *s*, respectively.

The attention-based deep neural network of DeepHINT has been implemented using the Keras library 2.0.8^2^, and four GTX 1080Ti GPUs have been used to accelerate the training and testing processes.

### 2.3 Baseline methods

#### 2.3.1 Score20

Implementation of Socre20 has been previously described [13]. Score20 depends on the local DNA sequence motif of the HIV integration sites (i.e., -10 bp to +10 bp) to discriminate positive samples from negative samples. Specifically, Score20 refers to the position weight matrix (PMW) corresponding to the log ratio of the base frequency in the positive samples versus that in matched random controls at each position. Note that the window length 20 bp has been shown to be the most effective parameter setting for HIV integration site prediction via this strategy [13].

#### 2.3.2 Random forest with genomic features

The random forest model is implemented following the protocol proposed in [14]. The input binary genomic features include ChIP-seq, DNase-seq and genomic annotation. ChIP-seq data, including H3K27Ac, H3K36me3, H3K4me1, H3K4me3, H3K9me3, RNA polymerase (Pol) II and CTCF as well as DNase-seq data, are collected from the ENCODE project [37, 38]. For the ChIP-seq data, we label the corresponding feature as 1 if the site of interest is located within 2 kb of the narrow peak center and 0 otherwise. To further characterize the relevant genomic features of HIV integration sites, we also include the transcription unit labels and the intron labels in the input features. We mainly use the RNA-seq data generated in the same HIV integration study [10] to obtain these labels. RNA-seq reads are aligned to the hg19 reference genome by HISAT2 and a transcriptome is assembled using Stringtie [39]. Only transcripts annotated in the Ensembl database [40] are kept and the expression levels of these genes are calculated as fragments per kb of exons per million (FPKM). Then the transcription unit feature is set to 1 if the site of interest lies in a transcript with FPKM higher than 1. Similarly, the intron label is set to 1 if the site of interest is located in the intron region of a transcript with FPKM higher than 1, according to the intron annotation from the Ensembl database [40]. The random forest model is implemented based on the Scikit-learn library [41], in which the maximum tree depth is set to 10 and tree number is set to 100. We further include the Score20 value as an additional feature in the random forest model to integrate local sequences as well as genomic features. A detailed list of data used is provided in Supplementary Table S2. Note that the same dataset is also used in the mechanistic analysis of DeepHINT prediction results (Section 3.2 and 3.3).

## 3 Results

### 3.1 DeepHINT accurately predicts HIV integration sites

We have performed extensive tests on known HIV integration sites in the HEK293T cell line [10] obtained from the Retrovirus Integration Database [15] and found that DeepHINT can significantly outperform the other state-of-the-art models in predicting HIV integration sites. To guarantee the data quality, only integration sites with equal to or more than five insertion counts were kept as positive samples. As the experimental determination of HIV integration sites involved a restriction enzyme (*MSEI*) digestion step that may lead to bias in the sequence context, matched random control sites were generated as negative samples following the same protocol described previously [10, 13, 14, 42]. More specifically, we first determined the genomic distances between all the positive samples to their nearest *MSEI* sites and then randomly sampled nine times more matched control sites that had the same distance distribution to their nearest *MSEI* sites as the negative samples. Note that the number of negative samples was set to be ten times as many as positive samples to reflect the natural imbalance of integration versus non-integration sites. To facilitate the training and evaluation of our model, the whole dataset was separated into strictly non-overlapping training and testing sets by chromosomes. Specifically, samples on chromosomes 1, 2, 3 were assigned to the test set, while the samples from the remaining chromosomes were used as the training set. Furthermore, for positive samples with genomic distances less than 1,000 bp, we only kept the one with the higher insertion count to avoid potential redundancy in the dataset. Overall, the aforementioned protocol resulted in 48,595 and 15,712 positive samples for training and testing, respectively, as well as a corresponding ten times larger set of negative samples. The final prediction performance was evaluated and reported based on the test data.

We first compared our method with a conventional position weight matrix (PWM) based method, namely Score20 [13], which directly calculates the consistency of the -10 to +10 bp window of a given site of interest with the consensus motif generated from training data. That is, Score20 mainly focuses on the local sequence motifs that favor HIV integrase binding. Expectedly, by efficiently integrating a much broader genome context of HIV integration, DeepHINT achieved a great improvement over Score20, with an increase of the area under the precision recall (AUPR) curve by 14.1% and the area under the receiver-operating characteristic (AUROC) curve by 5.5% (Fig. 3(a) and 3(b)). Note that in such a prediction task, the AUROC scores are prone to be boosted since the number of true negative samples is considerably larger than that of the false positive samples. Therefore, the precision-recall curves and the corresponding AUPR scores provide a more effective criteria for performance evaluation in our prediction problem.

**Figure 3:**
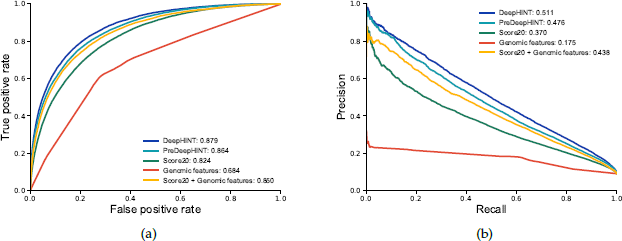
Prediction performance on the test dataset. (a-b) Comparison of prediction performance of DeepHINT with that of different baseline methods, in terms of (a) receiver-operating characteristic (ROC) curves and (b) precision recall (PR) curves, respectively. “preDeepHINT” denotes the DeepHINT framework without using the attention mechanism. The “Genomic features” mean the genomic profiles collected from the ENCODE project (see the main text for more details).

On the other hand, as also shown in the previous studies [10, 13, 14], the surrounding genomic features, e.g., chromatin accessibility, histone markers, transcription unit and intron annotation, also possess certain predictive information for detecting the retrovirus integration sites. Therefore, to test whether DeepHINT can sufficiently capture the genome context of HIV integration sites, we also compared its prediction performance to that of a random forest based model which explicitly required these additional surrounding genomic features as input, both with and without incorporating the Score20 values representing the local DNA sequence features (See Methods). In particular, the following experimentally measured genomic profiles were used as input to this random forest based model, including H3K27Ac, H3K36me3, H3K4me1, H3K4me3, H3K9me3, RNA polymerase (Pol) II, CTCF ChIP-seq data and DNase-seq data, as well as transcription unit and intron unit labels derived from RNA-seq data. We found that the random forest based model solely built on the above genomic features performed poorly (Fig. 3(a) and 3(b)), with an AUPR score of 17.5% and an AUROC score of 68.4%, indicating a necessity to effectively integrate various genomic context information in the modeling process. Intriguingly, although the integration of these additional genomic data did boost the prediction performance of Score20, DeepHINT still outperformed the random forest model by 7.3% in AUPR and 2.9% in AUROC (Fig. 3(a) and 3(b)). Note that such a comparison was biased to the random forest model as DeepHINT only took DNA sequence as input while the random forest model was fed with plenty of additional experimental data that have been shown to correlate with HIV integration. All these results demonstrated that the deep learning framework in DeepHINT can effectively learn the hidden feature representations encoded in the primary DNA sequences surrounding the HIV integration sites. Notably, we found that without using the attention mechanism, the prediction performance dropped significantly, with a decreased of 3.5% in AUPR and 1.5% in AUROC, which validated the contribution of attention mechanism to final prediction result of DeepHINT (Fig. 3(a) and 3(b)). We also conducted an additional conventional five-fold cross validation test over the whole dataset and observed a similar trend of the prediction results among different methods (Supplementary Figure 1).

### 3.2 The DeepHINT prediction scores associate with known genomic features

To further characterize how DeepHINT captures the various factors that influence HIV integration site selection, we also performed a series of statistical analyses to probe the possible associations between DeepHINT prediction scores and several known genomic features. To do this, we mainly compared the DeepHINT prediction scores in those regions enriched with individual genomic features, (i.e., all the genomic markers and annotations described in the previous section), to that of the background (i.e, randomly selected genomic positions). More specifically, for each feature identified by ChIP-seq and DNase-seq data, we randomly sampled 10,000 genomic loci within 1,000 bp of the centers of all narrow peaks. For the transcription unit and intron regions, we simply randomly sampled 10,000 genomic loci within them. All the analyses were performed on the test chromosomes, i.e., chromosome 1, 2 and 3, to eliminate the potential effect of overfitting in the training process.

As is shown in Fig. 4, the DeepHINT prediction scores displayed different levels of association with the various genomic features in our analyses, validating the general biological relevance of our approach. Expectedly, those sites near the PoI II binding sites or within the active transcription units (TUs) represented the most significant difference from the background regions (*P* < 10^−100^ for PoI II and *P* = 4.30 × 10^−65^ for TU, both by two-sided Wilcoxon rank-sum test), highlighting the effect of transcriptional activity in the selection of HIV integration sites [6, 9]. Consistently, the histone markers labeling actively transcribed regions also exhibited significantly higher HIV integration probability. In particular, the active promoter marker H3K4me3 and the general active transcription marker H3K27ac both showed a higher DeepHINT prediction score (*P* = 4.47 × 10^−16^ for H3K4me3 and *P* = 9.03 × 10^−11^ for H3K27ac, both by two-sided Wilcoxon rank-sum test). The same phenomenon was also observed for H3K36me3 (*P* = 6.34 × 10^−14^ by two-sided Wilcoxon rank-sum test), which has been previously shown to be associated with both active gene transcription and HIV integration [45, 46], probably through the direct binding of the integration factor LEDGF/p75 [47]. In contrast, those histone markers indicating active enhancers (i.e., H3K4me1) and silent gene regions (i.e., H3K9me3) did not show any significant difference from the background (*P >* 0.01 by two-sided Wilcoxon rank-sum test for both), suggesting a less important role of these factors in HIV integration. Interestingly, DNase hypersensitive regions (i.e., DNase-seq peaks) presented a significant correlation with the DeepHINT prediction score (*P* = 2.12 × 10^−34^ by two-sided Wilcoxon rank-sum test), indicating that the DNA accessibility also plays an important role for HIV integration, possibly through the binding of some regulatory factors. In addition, consistent with the previous studies [10, 48–50], we also noticed a significant enrichment of the DeepHINT prediction scores in the intron regions and near CTCF binding sites (*P* = 1.98 × 10^−11^ for intron regions and *P* = 1.52 × 10^−14^ for CTCF binding regions, both by two-sided Wilcoxon rank-sum test), further demonstrating the broad range of information captured by the DeepHINT model. Note that a negative control test on an additional sets of randomly-selected genomic sites was also carried out to support the biological relevance of our analyses.

**Figure 4:**
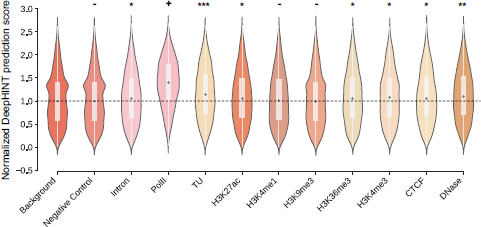
Comprehensive statistical analyses between the DeepHINT prediction scores and various known genomic features. Data labels: negative control (randomly-selected 10,000 codon sites), Pol II (RNA polymerase II), TU (active transcription unit). *: 5 × 10^−25^ < *P* < 1 × 10^−2^; **: 5 × 10^−50^ < *P* ≤ 5 × 10^−25^; ***: 5 × 10^−100^ < *P* ≤ 5 × 10^−50^; +: *P* ≤ 5 × 10^−100^; −: *P >* 0.01; two-sided Wilcoxon rank-sum test.

### 3.3 DeepHINT indicates important sequence positions for predicting HIV integration sites

The involvement of attention mechanism opened up the black box of deep learning and further enabled us to probe the derived attention map for each sample. We hypothesize that the positions with larger attention values are more likely to associate with the sequence determinants of HIV integration site selection. Therefore, we were particularly interested in the *attention intensive regions* (which were defined as those the genomic positions possessing the highest 5% attention values of an input sequence) and digging out how they can reflect the underlying biological mechanisms of HIV integration. Note that due to the convolution (whose kernel size was set to 6) and max pooling (whose pool size was set to 3) operations, each position in the attention map (also called the *attention map index*) represented a 8-bp region consisting of three continuous convolution kernels.

As a first attempt, we performed a close-up inspection for the distribution of the attention intensive regions near the integration sites for all positive and negative samples in the test dataset (Fig. 5(a)). Intriguingly, we observed a distinct pattern in the distributions of attention intensive regions between positive and negative samples. In particular, the distribution of attention intensive regions in positive samples showed a clear peak-valley-peak pattern near the integration sites, which, on the other hand, was not observed in that of negative samples. Such a discrepancy indicated that the attention map derived from our deep learning model was able to reflect the sequence specificity pattern in the genomic context of HIV integration. Next, we further compared the above pattern derived from the attention intensive regions with the local consensus sequence motif obtained from the Score20 method and found that the shape of the attention profile aligned well with the conserveness of each nucleotide in the Score20 motif (Fig. 5(a) and 5(b)). In particular, the sequence windows corresponding to attention map indexes 1 and 4, which included the most conservative G11 and C15 nucleotides in the Score20 consensus motif, showed the highest attention scores. On the other hand, the “attention valley” in the observed attention profile also matched the non-conservative region in the Score20 motif (i.e., sequence windows corresponding to indexes 2 and 3).

**Figure 5:**
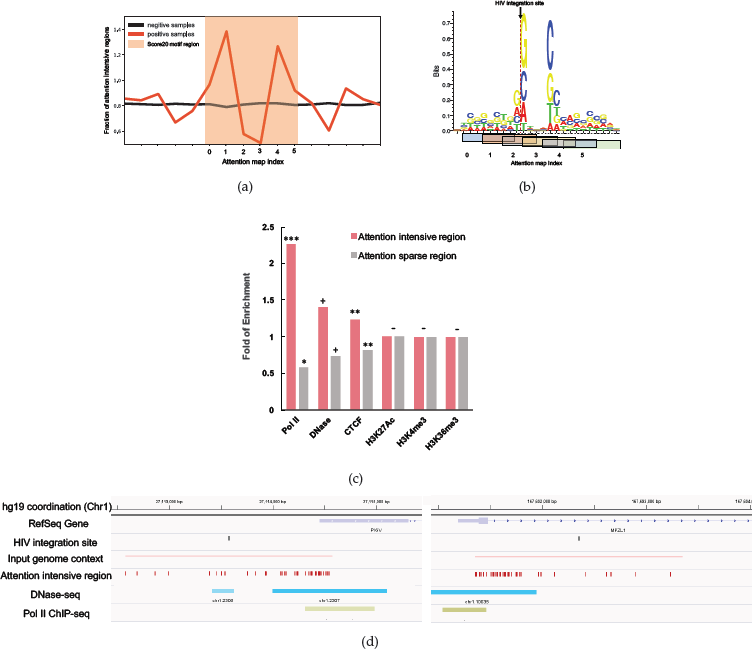
Attention intensive regions indicate important local features of the predicted HIV integration sites. (a) A local view of the distribution of attention intensive regions near the integration sites for both positive samples (red) and the matched random control sites of negative samples (black) in the test dataset. The region overlapping with the Score20 consensus motif is highlighted (orange). Fractions of attention intensive regions were averaged among all the samples and normalized to the mean of all positions. Note that due to the convolution (whose kernel size was set to 6) and max pooling (whose pool size was set to 3) operations, each position in the attention map (termed as the attention map index) represented a 8-bp region consisting of three continuous convolution kernels. (b) The Score20 consensus motif after being aligned with the average attention profile. The attention window index corresponding to (a) is shown as x-axis and the sequence windows corresponding to each individual indexes are labeled below. The DNA WebLogo is visualized using Seq2Logo [43]. (c) The levels of enrichment or deleption of different genomic features for both attention intensive regions (with the highest 5% attention, pink) and attention sparse regions (with the lowest 5% attention, gray). Fold of enrichment was calculated by using 5% randomly selected regions as baseline. *: 5 × 10^−5^ < *P* < 1 × 10^−3^; **: 5 × 10^−10^ < *P* ≤ 5 × 10^−5^; ***:5 × 10^−40^ < *P* ≤ 5 × 10^−10^; +: *P* ≤ 5 × 10^−40^; −: *P >* 0.05; Chi-square test. (d) Two representative examples illustrating the enrichment of attention intensive regions in PoI II binding and DNase hypersensitive regions. The visualization was conducted using the IGV browser [44].

Given the great improvement in the prediction performance of DeepHINT over the Score20 method (Fig. 3(a) and 3(b)), we were interested in how DeepHINT learns the sequence features beyond the Score20 region. Therefore, we further examined the associations of attention intensive regions with diverse genomic profiles and showed that the important genomic positions indicated by the attention mechanism can provide more mechanistic insights into the integration process. In particular, we performed a series of statistical analyses on the levels of enrichment or depletion of different genomic features in the attention intensive regions and the attention sparse regions (defined by the regions with lowest 5% attention) (Fig. 5(c)). Here we mainly focused on those factors that were shown to significantly associate with the DeepHINT prediction scores (Fig. 4). Expectedly, PoI II binding sites exhibited a significant association with the attention intensive regions, showing a 3.88 fold of enrichment in the attention intensive regions compared to that of the attention sparse regions. In addition, the DNase hypersensitive regions displayed the most statistically significant enrichment and depletion in the attention intensive regions and sparse regions, respectively (*P* < 10^−40^ by Chi-square test), further supporting a potential role of the binding of regulatory factors in the chromatin accessible regions for HIV integration. However, for the peaks of histone markers, i.e., H3K36me3, H3K27ac and H3K4me3, which commonly spread across the sequence context of HIV integration, in principle we cannot detect large signal difference between attention intensive and sparse regions (Fig. 5(c)), which also indicated the necessity to examine the more localized genomic features, e.g., sequence motifs, that may be easier to be captured by the attention mechanism. Two representative examples of the enriched attention near the Pol II binding regions and DNase hypersensitive regions can be found in (Fig. 5(d)).

### 3.4 DeepHINT highlights the sequence features for HIV integration site selection

The clear association between the derived attention intensive regions and chromatin accessibility suggested the potential capacity of DeepHINT to reveal regulatory factors that may play an important role in the selection of HIV binding sites. To further exploit the specific sequence features captured by our attention mechanism, we also conducted a systematic survey on the sequence enrichment in those attention intensive regions. More specifically, we extracted all the 8-bp sequences in the attention intensive regions and calculated the enrichment of the binding motifs of known mammalian DNA binding proteins using HOMER [52].

Intriguingly, we identified two families of important regulatory factors, i.e., bHLH transcription factors and zinc finger proteins, whose binding sites showed significant enrichment in the attention intensive regions (Fig. 6). The binding of bHLH transcription factors recognizes E box (i.e., CACGTG), a highly conserved eukaryotes cis-regulatory DNA element that plays a wide range of regulatory roles in controlling gene activities [53]. In retrovirus, the presence of E box motifs in long terminal repeats (LTRs) have been shown to associate with the modulation of gene expression and maintenance of virus latency [54], probably by repressing the expression of viral proteins [55, 56]. However, how the E box element in human genome sequence participates in the regulation of HIV latency still remains unclear. Similarly, the zinc finger protein ZFX has also been shown to be involved in the modulation of LTR mediated HIV transcription [57]. Here, our results may provide two possible directions to further explore the cis-regulatory factors in human genome that may contribute the HIV integration site selection.

**Figure 6:**
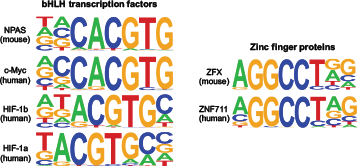
Attention intensive regions are enriched with the binding motifs of bHLH and zinc finger proteins. The known mammalian motifs obtained from TRANSFAC [51] that were significantly enriched in attention intensive regions are shown. All the motifs showed a Benjamini q-value < 10^−4^ as determined by HOMER [52].

## 4 Discussion

Despite the long-lasting experimental and computational effort devoted to study HIV integration [10, 13, 14, 58], our current understanding of the mechanistic implications of HIV genome integration still remains largely limited. In this study, instead of focusing on the tedious hand-crafted feature engineering, we developed an attention-based deep learning framework, namely DeepHINT, to automatically learn the contextual sequence features of HIV integration, and precisely predict the integration sites. In addition to boosting the prediction performance, the attentions map derived by DeepHINT can explicitly decode how the deep learning model recognized highly relevant sequence features at different positions for final prediction, enabling researchers to better understand the underlying mechanism of the HIV integration. Extensive tests validated that DeepHINT greatly outperformed the conventional prediction models and achieved the state-of-the-art performance in identifying HIV integration sites. In-depth analyses also demonstrated the biological relevance and several application potentials of DeepHINT for future mechanistic studies of HIV integration.

The current study demonstrates the usage of deep learning, especially with the attention mechanism, for predicting and analyzing HIV integration sites. Admittedly, the further exploration of the underlying mechanism of HIV integration also relies on the generation of high-quality HIV integration sites, especially in human patients [4, 5]. Nevertheless, the introduction of effective machine learning techniques, e.g., using transfer learning to transfer the cell line knowledge to patient samples would also be an interesting future direction to explore. As integration-associated virus latency is attracting more and more research interest [59], we believe that our DeepHINT framework together with more emerging experimental data [15] and improved experimental techniques [60] will offer more useful insights into the studies of HIV integration in the genome.

## Footnotes

1 https://github.com/maxpumperla/hyperas/

2 https://keras.io/

